# Influence of gut microbiota on oral drug absorption and metabolism

**DOI:** 10.1101/2025.02.16.638186

**Authors:** Douglas Nelson, Vaishnavi Veerareddy, Purna C Kashyap, Xiaojiao Tang, Krishna R. Kalari, Karunya K. Kandimalla

**Author notes:** Corresponding author: Karunya K. Kandimalla, Department of Pharmaceutics and Brain Barriers Research Center, College of Pharmacy, University of Minnesota, Minneapolis, MN, 55455, USA. **Conflict of interest statement** All other authors declared no competing interests for this work.

## Abstract

Gut bacteria influence host intestinal phenotype in ways that impact drug absorption and metabolism. By comparing colonic tissue from germ-free mice to that of mice colonized with human microbiota, this study evaluates microbiota-driven differences in gene expression, mucosal permeability, and P-gp efflux capacity. Transcriptomic analysis revealed upregulation of genes coding ATP-binding cassette drug transporters P-gp (Abcb1a, Abcb1b), BCRP (Abcg2), and MRP3 (Abcc3), along with increased expression of solute carriers MCT1 (Slc16a1) and OCTN2 (Slc22a5). Immunohistochemistry indicated greater P-gp expression and apical localization in humanized mice. No relevant gene expression changes were observed in human homologs of drug-metabolizing cytochrome P450 enzymes, though cytochrome P450 oxidoreductase (Por) upregulation suggests increased cytochrome activity. Other phase I drug-metabolizing enzymes, including multiple homologs of human carboxylesterase 2 (CES2) and several reductases, were upregulated with high significance. Regarding phase II metabolism, genes encoding most UDP-glucuronosyltransferases and glutathione S-transferases were upregulated, along with the enzymes responsible for synthesizing their corresponding co-substrates, UDP-glucuronic acid and glutathione. Small intestinal mucosal explants demonstrated higher permeability to ^14^C-labeled polyethylene glycol 4000 (^14^C-PEG4000) in germ-free mice than in humanized mice. The ^14^C-PEG4000 permeability differences are in agreement with changes in paracellular tight junctional proteins such as claudins, zonula occludens, and myosin II-related genes. These results demonstrate the broader impact of gut microbiome on oral drug absorption and metabolism and implicate the contributions of gut microbiome to individual variations in drug efficacy and toxicity.

## INTRODUCTION

The human gut microbiome influences drug bioavailability through multiple mechanisms. In addition to directly supplying drug-metabolizing enzymes, gut bacteria regulate the expression of intestinal enzymes, transporters, tight junctions, and other molecules affecting oral drug disposition. By comparing germ-free mice to those colonized with human fecal microbiota, this study is aimed at investigating microbiota-induced phenotypic changes relevant to drug absorption and metabolism.

The extent to which intestinal phenotype influences oral drug absorption depends on a drug’s physical properties—primarily permeability, solubility, and lipophilicity—which determine its absorption potential and susceptibility to enzymatic transformation^1^. High-permeability, high-solubility drugs, such as acetaminophen, pass rapidly through the intestinal epithelium and are only modestly impacted by transporters and metabolizing enzymes^2^. Drugs that show low permeability and high solubility, such as metformin, often require active transport for absorption and are more susceptible to changes in the expression of drug transporters and intestinal metabolism^3^. On the other hand, highly permeable but poorly soluble lipophilic drugs, such as ketoconazole, exhibit low intestinal absorption as they are effluxed by transporters like P-gp^4^. In cases of small hydrophilic molecules with low permeability, such as mannitol, absorption depends almost entirely on paracellular transport through tight junctions^5^.

Two primary enzymatic pathways that drive the metabolism of a majority of drugs are considered in this paper: Phase 1 metabolism, which modifies drugs via oxidation (primarily by cytochrome P450 enzymes), reduction, or hydrolysis; and phase 2 metabolism, which forms drug metabolites by conjugation to increase their solubility and facilitate excretion. While phase 2 metabolism often follows phase 1, some drugs undergo only one of these processes^6^.

Microbiota density increases by roughly ten orders of magnitude from the proximal small intestine to the colon, which implies a progressively stronger microbial influence on gene expression and metabolism^7^. This variation affects both direct microbial metabolism of drugs and microbiota-mediated regulation of intestinal enzymes, transporters, and tight junctional integrity^8^. Microbial metabolites play an important role in this regulation by modifying the intestinal environment and influencing host gene expression^8^.

The small intestine is the primary site of drug absorption, with the duodenum, jejunum, and ileum accounting for most systemically absorbed drugs^9^. The relative contribution of each small intestinal segment varies by drug. For example, drugs with high solubility and high permeability drugs can be largely absorbed in the duodenum before reaching other sections, while drugs with lower solubility tend to be absorbed more efficiently in the jejunum and ileum due to their extensive microvillar surface area^10^. While most systemically active drugs are absorbed in the small intestine, the colon remains relevant for formulations targeting local conditions such as inflammatory bowel disease. In addition, some systemically active drugs, such as acetylsalicylic acid, exhibit measurable, albeit minor, colonic absorption^11^. Despite its limited role in drug absorption, the colon is the primary site where microbiota regulate intestinal phenotype, making it a reasonable starting point for studying microbiota-driven changes in transporters, enzymes, and barrier properties.

## METHODS

### Mouse Husbandry

All animal studies were approved by the Mayo Clinic Institutional Animal Care and Use Committee and conducted in accordance with regulatory guidelines. Mice were housed under a 12-hour light-dark cycle and monitored daily for any signs of physical stress or behavioral changes. If distress was observed, mice were euthanized per protocol. Germ-free (GF) Swiss Webster mice were obtained from the Mayo Clinic Germ-Free Facility, and humanized mice were generated from GF Swiss Webster mice. Both GF and humanized mice were housed in flexible film vinyl isolators (Class Biologically Clean, Madison, WI) within the Mayo Clinic Germ-Free Facility, where they were kept in open-top cages with autoclaved Sani-Chips bedding, an autoclaved diet (Purina Lab Diet, 5K67, Collins Feed and Seed Center, Rochester, MN), and autoclaved nanopure water. Bedding and feed were changed weekly or sooner if needed. GF status was confirmed prior to experiments through two consecutive cultures of fecal pellets, feed, and bedding on brain heart infusion (BHI), Sabouraud dextrose, and nutrient media, under both anaerobic and aerobic conditions. Additionally, PCR analysis of 16S rRNA genes was performed using universal and Turicibacter-specific primers on fecal DNA. Humanized mice were generated by oral gavage of stool from a healthy human donor into GF mice at 4–5 weeks of age. The stool suspension was prepared by mixing equal volumes of stool and sterile pre-reduced PBS inside an anaerobic chamber (Coy Laboratory Products, Grass Lake, MI). The sealed vial was then removed from the chamber and transferred to GF isolators. Each mouse received 300 μL of the stool suspension via gavage and was allowed 4–5 weeks for microbiota adaptation to the mouse gut.

### Transcriptomics: Library Preparation, Sequencing and Alignment

Messenger RNA was extracted from mouse samples for RNA-Seq library preparation following the instructions of the Illumina TruSeq RNA Library Prep Kit v2. Sequencing was performed on an Illumina High Seq-3000 by the Mayo Clinic Medical Genome Facility with 101bp paired end reads. Sequencing data was collected with Illumina HCS v3.3.52. Base-calling was performed with Illumina RTA v2.7.3 and converted to fastq with Illumina BCL2FASTQ v2.17.1.14. The FASTQ files were processed using the Mayo Clinic in-house pipe-line MAP-RSeq v.2.1.1 workflow^12^. Sequences were aligned with TopHat (2.1.0) and Bowtie (2.2.3) against the mouse genome build mm10. Gene counts were quantified using featureCounts from Subread package 1.4.6-p5 with mm10 mouse gene annotation^13^. The resulting counts matrix of with 43,346 mus musculus ensemble genes is included in the supplementary material.

### Transcriptomics: Analysis

Mouse Genome Informatics “complete list of human and mouse homologs” downloaded from https://www.informatics.jax.org/homology.shtml. A human homolog database was constructed with 26,281 homologous pairs exhibiting one-to-many relationships in both directions. All statements regarding human-mouse homologs in this study are based on these homology assignments. Low expression filtering, TMM normalization, dispersion estimation and differential expression testing with Benjamini Hochberg multiple hypothesis correction was performed with edgeR 4.0.16, using the classic pipeline with default parameters. A total of 19,863 genes remained after low-expression filtering. Relative gene abundance was estimated by RPKM (Reads Per Kilobase per Million mapped reads) with TMM normalization using the edgeR RPKM function, essentially providing gene-length normalized model-fitted counts. The percentile distribution was based on all samples. Results of normalization, differential expression and relative abundance estimation are included in supplementary files. Over-representation testing of differential expressions in phase 2 enzyme families was calculated using the hypergeometric test (one-sided Fisher’s exact).

### Organoid studies

Three-dimensional colonic organoids were prepared by isolating the proximal colon segment from 8-to 12-week-old germ-free and humanized mice. Intestinal crypts were centrifuged in cold DMEM/F12 medium (Life technologies, Carlsbad, California), and the pellet was re-suspended in the IntestiCult™ Organoid Growth Medium (StemCell Technologies, Vancouver, Canada) on ice. Reduced Matrigel (1:1 ratio, Corning) was then added to the crypt suspension and mixed thoroughly. An 80 µl aliquot of this suspension was added to a pre-warmed 24-well plate, creating a dome-like structure, which was incubated at 37°C for 10 minutes before adding medium supplemented with penicillin/streptomycin. The medium was changed every 3–4 days depending on organoid growth. After 7 days, organoids were passaged by re-dispersing them with pipette in gentle Cell Dissociation Reagent (StemCell Technologies, Vancouver, Canada) at 37°C for 15 min. Dispersed organoids were diluted with DMEM/F12 and pelleted by centrifuging (290×g) for 5 minutes at 4 °C. The pellet was then resuspended in organoid growth medium and plated on coverslip-bottom dishes.

### Organoid efflux and cell viability studies

Efflux studies were conducted to investigate P-gp functionality in colonic organoids using calcein-AM, a P-gp substrate. Organoids growing on coverslips were incubated in medium supplemented with calcein-AM (0.5 µM) for 60 minutes at 37 °C in the control group. The treatment group was pretreated with the P-gp inhibitor cyclosporin A (5 µM) for 30 minutes before adding calcein-AM. Live images were taken under 37 °C and 5% CO₂ using a Nikon TE2000-S inverted microscope (Chiyoda-ku, Tokyo 100-8331, Japan) equipped with a Nikon FITC HQ filter and a Zeiss Axio Observer Z1 spinning disk laser confocal microscope equipped with a Photometrics QuantEM 512C EMCCD camera. All images were processed in Fiji software (v1.54f), where they were subjected to automated background subtraction using a rolling ball radius of 50 pixels.

### Immunohistochemistry

Colonic tissue was collected from mice and fixed with 4% paraformaldehyde in 1X Dulbecco’s Phosphate-Buffered Saline (PBS) at 4 °C for 4 h. Tissue was then dehydrated in a gradient of solutions with various concentrations of sucrose [10% (w/v), 20% (w/v), and 30% (w/v)] over 3 days at 4 °C. After dehydration, the tissue was embedded in the Tissue-Tek O.C.T. compound (Sakura Finetek). The embedded tissue was sectioned into 8 µm slices using a cryostat (Leica Biosystems CM3050 S Research Cryostat) and mounted on slides.^14^ Endogenous peroxidase activity was blocked by incubating the sections in 3% hydrogen peroxide in distilled water. Subsequently, sections were blocked using goat serum (1:10 in 0.02 M PBS) for 20 minutes at room temperature. The blocked sections were incubated overnight at 4 °C with a P-gp antibody (anti-rabbit IgG, RP1034, Boster Biotechnology, Pleasanton, CA, 1:200). The next day, tissue was washed three times with PBS, followed by incubation with a biotinylated goat anti-rabbit IgG secondary antibody (1:200) at 37 °C for 30 minutes. Tertiary antibody staining for P-gp was performed using Cy3-conjugated streptavidin (SABC-Cy3, 1:200) at 37 °C for 30 minutes in the dark. Finally, nuclei were stained by incubating the tissue with 4′,6-diamidino-2-phenylindole (DAPI, 1:5000 in 0.02 M PBS). Immunofluorescence was detected using a Leica inverted fluorescence microscope (Leica Microsystems, Buffalo Grove, IL) and captured using Leica Application Suite X (LASX) software. Images from two germ-free and three humanized colon samples were analyzed using FIJI software (v1.54f). After threshold background removal, mean intensity of each image sample was calculated. Additionally, mean intensity of 50 manually selected points along the apical surface was calculated. Then the means of germ-free and humanized sample intensity values were compared by two-tailed unpaired t-test.

### Permeability studies

Gastrointestinal mucosal permeability of small intestinal tissue from germ-free and humanized mice was assessed in vitro using an Ussing chamber. After collecting epithelial tissue from the small intestine, it was opened along the mesenteric border and washed in ice-cold Kreb’s solution. The mucosa-submucosa epithelial sheet was mounted onto the Ussing chamber cassette such that the serosal side faced up, and the muscle layer was peeled away. The basolateral and apical sides of the chambers contained glucose-containing Krebs solution and Krebs-mannitol solution, respectively. The ^14^C-PEG 4000 (0.25 µCi) was added to the mucosal side, and the samples were collected from the serosal side at 1, 5, 10, 15, 30, 60, and 90 min to determine the mucosal-serosal permeability as described previously^14,15^. Radioactivity transported to the serosal compartment was assessed in the beta-counter. The flux was determined from the slope of the linear portion of cumulative amount in the serosal compartment versus time plot as described previously and the permeability was determined by normalizing flux with the initial donor concentration of ^14^C-PEG 4000. A two-tailed unpaired t-test was used to compare humanized and germ-free samples.

## RESULTS

### Transcriptomic overview and differential gene expression

The principal component analysis (PCA) plot (Figure 1) illustrates that germ-free and humanized mice represent distinct transcriptomic profiles. These groups form clusters that are well-separated by the first principal component, representing most of the total sample variation in gene expression. Both principal components contribute to variation among humanized mice, which show higher overall dispersion than germ-free mice. Comparing the expression profiles of 19,863 genes in humanized mouse colon to germ-free mouse colon resulted in 3,826 differentially expressed genes with false discovery rate (FDR) corrected p-values less than 0.01. The number of differentially expressed genes at various thresholds of statistical significance and magnitude is presented in Table 1.

**Figure 1:**
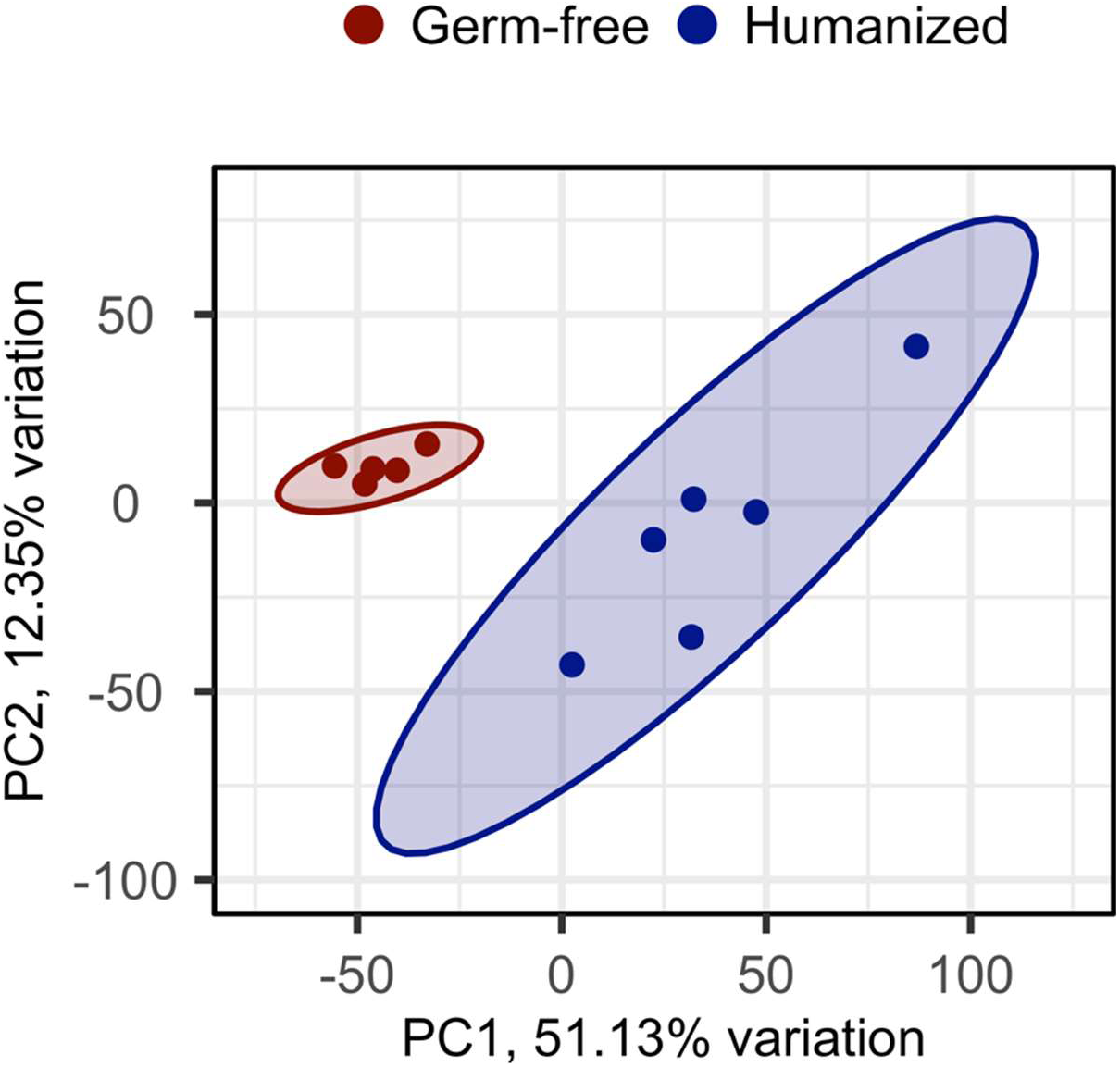
Principal component analysis of RNA-Seq results for colons of five germ-free (red) and six humanized (blue) mice. Ellipses denote the borders of the 95% confidence intervals for each group. First principal component (PC1) represents 51% of the overall genewise variation and largely accounts for the variance among germ-free samples. Both principal components contribute to the variation among humanized mice, which exhibit greater overall dispersion compared to germ-free mice.

**Table 1:**
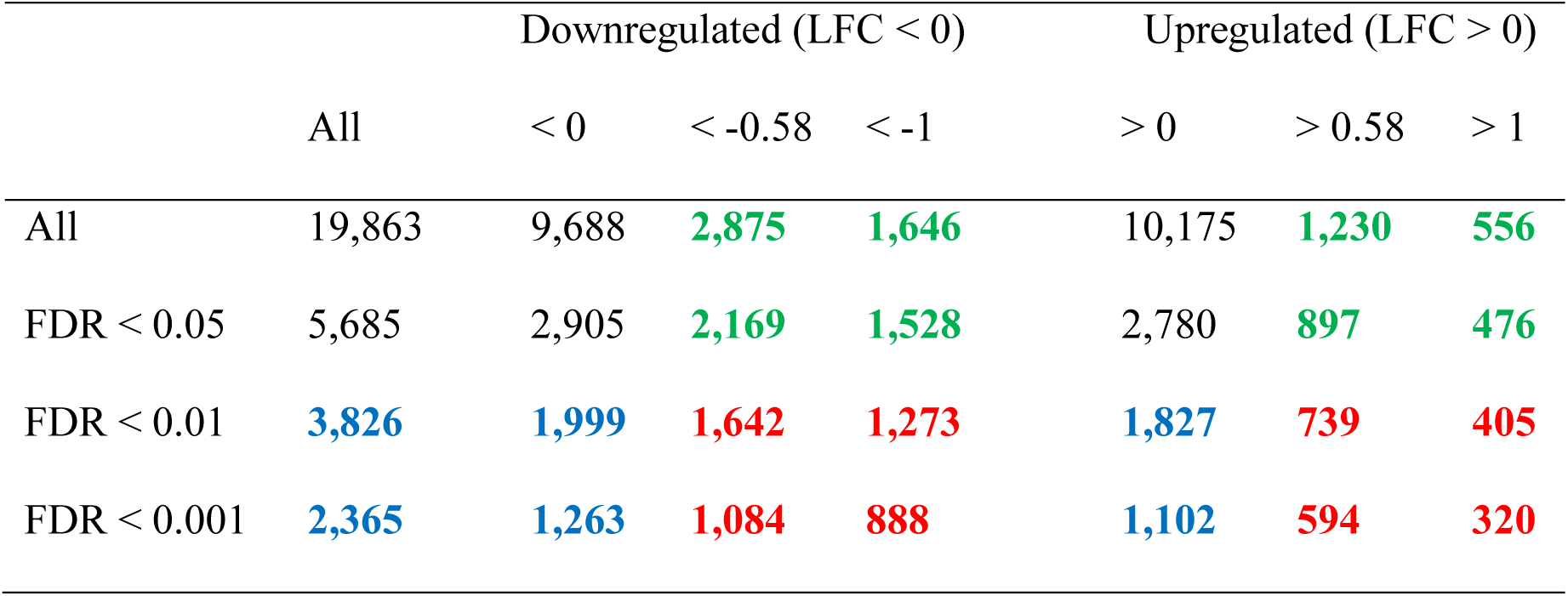
Result of differential expression analysis showing the number of genes corresponding to various thresholds of statistical significance and directed magnitude. As in the volcano plots, blue denotes genes with a false discovery rate of less than 0.01, green represents genes with log2-fold change absolute magnitude of greater than 0.58, and red indicates genes with both FDR < 0.01 and absolute log2-fold changes greater than 0.58, corresponding to fold-changes of 2/3 and 3/2.

### Immune response and extracellular matrix

The introduction of human microbiota resulted in a general increase in colonic gene expression related to adaptive immunity. This includes constant and variable chains that comprise IgG and IgA immunoglobulins; the polymeric immunoglobulin receptor (Pigr), crucial for epithelial secretion of immunoglobulins, as well as H2-Q1 and H2-Q2, which are mouse homologs of human HLA-E. Although immunoglobulin chains exhibited the largest mean fold changes, p-values were relatively high because of high sample dispersion. On the other hand, genes central to epithelial innate immunity were moderately upregulated with relatively uniform inter-sample expression and high statistical significance. This includes the reactive oxygen species (ROS)-producing nitric oxide synthase (NOS2) and dual oxidase complex genes (Duox1, Duox2, Duoxa1, Duoxa2). Log2-fold changes and FDR p-values illustrating this immune response are given in Figure 2.

**Figure 2:**
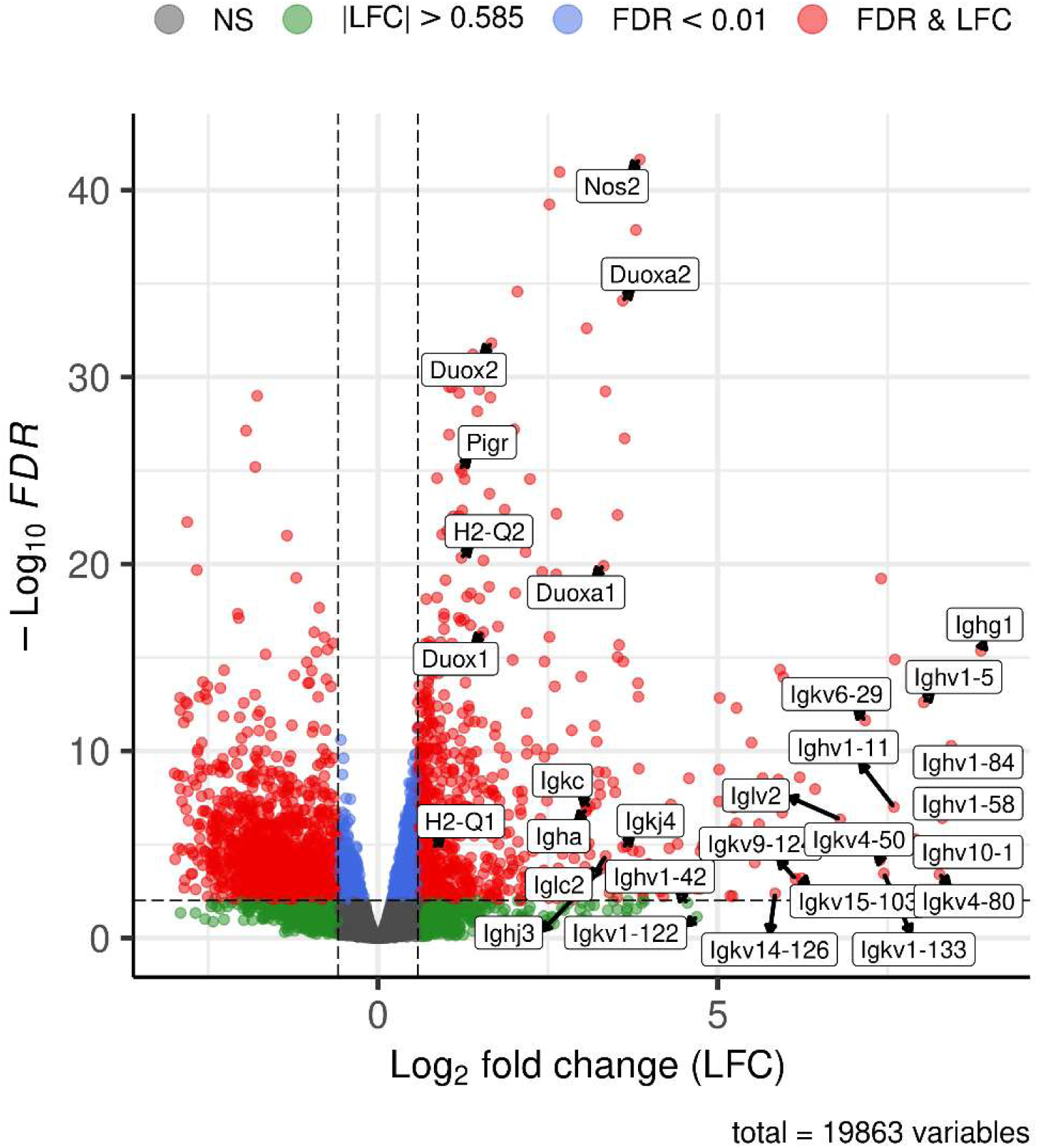
Volcano plot shows upregulation of certain genes related to innate and adaptive immunity in mice with humanized gut microbiota. Genes coding constant and variable chains that comprise IgG and IgA immunoglobulins are among the most upregulated in the entire gene set, with relatively high log-fold changes but also greater inter-sample variability, as indicated by their p-values. Other key genes related to adaptive immune response include the polymeric immunoglobulin receptor (Pigr), crucial for epithelial secretion of immunoglobulins, as well as H2-Q1 and H2-Q2, which are mouse homologs of human histocompatibility complex, class I, E (HLA-E). The indicators of innate immune response involving reactive oxygen species production are among the most statistically significant upregulated genes, including all the components of the dual oxidase complex (Duox1, Duox2, Duoxa1, Duoxa2) and Nitric Oxide Synthase 2 (Nos2), which is also involved in the maintenance of vascular tone. Nos2 was among the top 10 differentially expressed genes by statistical significance with FDR of 2.3*10-42.

In contrast to the immune response, Figure 3 shows that all the major constituents of the extracellular matrix (ECM), along with genes responsible for regulating ECM remodeling, are uniformly downregulated in humanized mouse colon. This includes genes encoding intestinal collagens in the basement membrane (collagen IV, VI, VIII, and XXIII) and the interstitial matrix (collagens I, III, V, XXII, and XVIII).^16^ Other ECM protein coding genes exhibiting similar differential expression included several key intestinal laminin subunits, as well as fibulins (Fbln1, Fbln2), nidogens (Nid1, Nid2), fibronectin (Fn1), decorin (Dcn), perlecan (Hspg2), agrin (Agrn), and osteonectin (SPARC). Additionally, humanized mouse colon also exhibited lower expression of genes related to ECM degradation affecting the ECM turnover rate, including metalloproteases (Mmp7, Mmp9, Mmp2) and the two metalloprotease inhibitors with biologically relevant expression (Timp2, Timp3).

**Figure 3:**
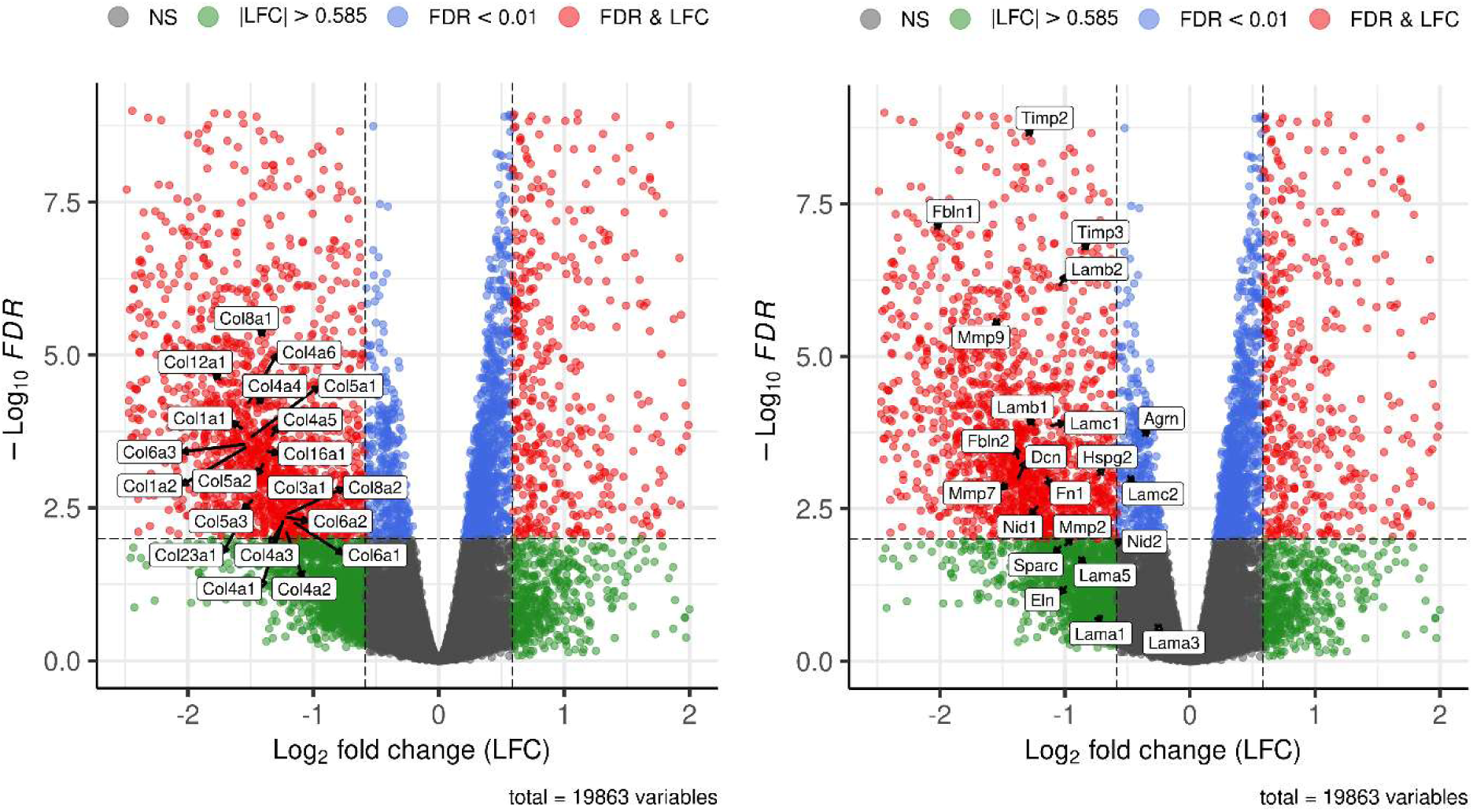
Genes coding all the major extracellular matrix components were less expressed in humanized mouse colon compared to those of germ-free mice. This includes all genes coding the intestinal collagens in the basement membrane (Collagen IV, VI, VIII and XXIII) and the interstitial matrix (Collagens I, III, V, XII, and XVIII). Other ECM protein coding genes exhibiting similar differential expression included several chains that comprise the major intestinal laminins, as well as fibulins (Fbln1, Fbln2), nidogens (Nid1, Nid2), fibronectin (Fn1), decorin (Dcn), perlecan (Hspg2), agrin (Agrn), and osteonectin (SPARC). Additionally, humanized mice colon also showed lower expression of genes related to ECM degradation affecting the ECM turnover rate, including metaloproteases (Mmp7, Mmp9, Mmp2) and the two metalloprotease inhibitors with biologically relevant expression (Timp2, Timp3).

Despite being downregulated, collagen and laminin genes show higher inter-sample variability in humanized samples. This is illustrated in Figure 4, which includes the genes coding the predominant collagen and laminin isoforms (collagen IV, Laminin-111, and laminin-511 of the basement membrane, as well as collagen I and collagen III of the interstitial matrix).

**Figure 4:**
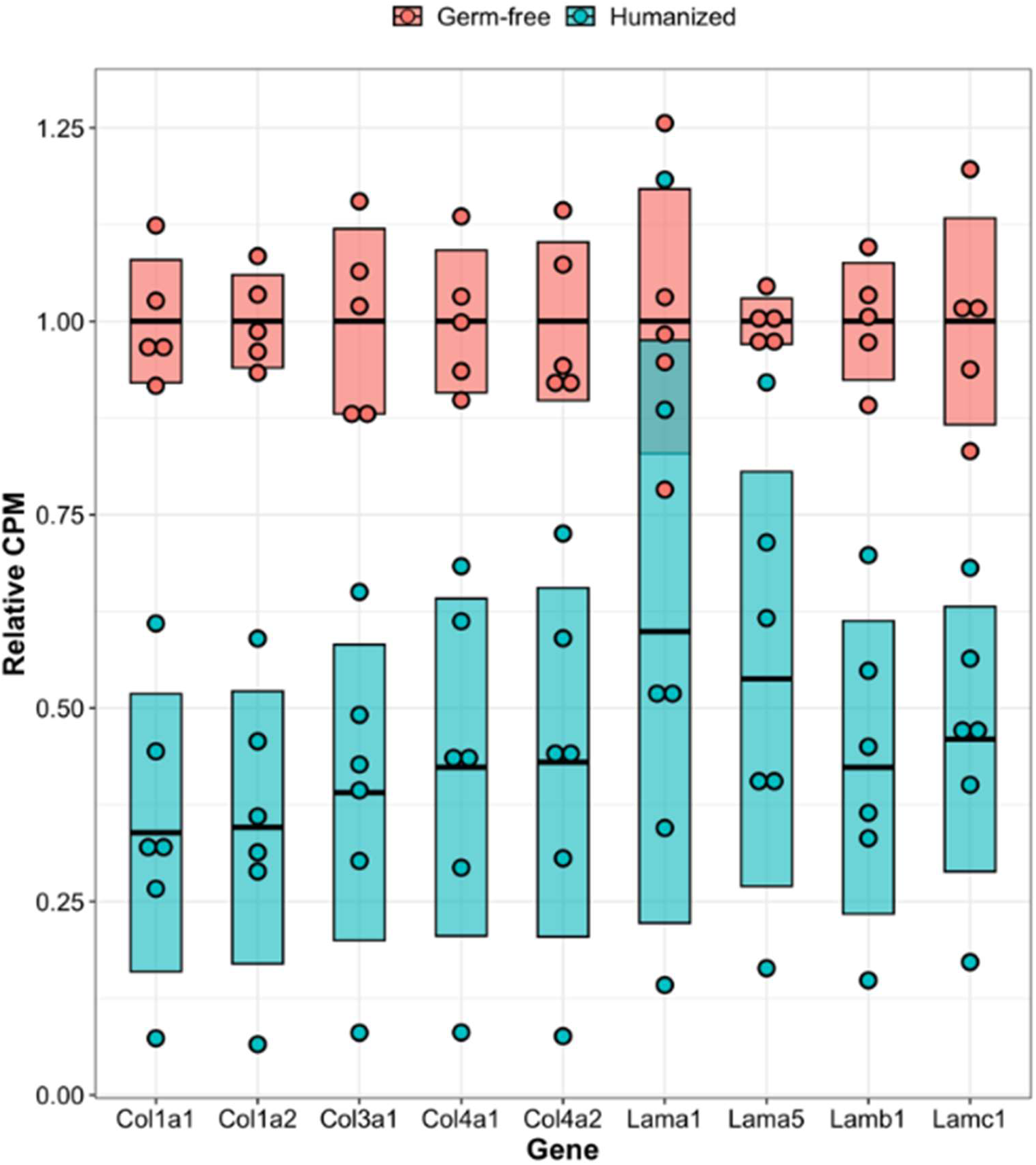
mRNA counts for key extracellular matrix coding genes relative to mean germ-free counts. Included are genes coding the predominant proteins that comprise the extracellular matrix: collagen IV, Laminin-111, laminin-511 of the basement membrane, as well as collagen I and collagen III of the interstitial matrix. Plot shows greater absolute dispersion among humanized mice, despite lower counts, suggesting ECM may be affected by variable composition of microbiota shaped by heterogenous immune response.

### Tight junctions and water channels

Table 2 presents the expression analysis of genes encoding proteins involved in tight junction structure (claudins, occludin, and zonula occludens proteins) and actomyosin contractility (myosin II and myosin light chain kinase, regulating myosin phosphorylation). Zonula occludens 3 (Tjp3), claudin 23 (Cldn23), and claudin 4 (Cldn4) were upregulated in humanized mouse samples. In particular, the changes in these two claudins were highly significant (FDR = 2.8 × 10^-23^ and 2.8 × 10^-11^). All three differentially expressed genes were highly expressed in the top RPKM decile, with Tjp3 being the most abundant zonula occludens.

**Table 2:** Differential expression and relative abundance of claudin, occludin and zonula occluden genes coding tight junction proteins, in mouse colon. LFC is the log2-fold change comparing colons with humanized gut microbes with those from germ-free mice. RPKM represents the TMM normalized mean RPKM values in the humanized samples and “%” represents the percentile in the humanized RPKM distribution. Listed here are all claudin, occluding, zonula occluden and myosin II-related genes with RPKM > 4. Genes coding claudin23 and claudin 4 were upregulated with very high significance.

| Gene | LFC | FDR | RPKM | % |
| --- | --- | --- | --- | --- |
| Tight Junctions |  |  |  |  |
| **Cldn23** | **1.11** | **2.8E-23** | **50** | **87** |
| **Cldn4** | **1.21** | **2.8E-11** | **34** | **79** |
| **Tjp3** | **0.32** | **5.6E-03** | **80** | **96** |
| Tjp2 | 0.14 | 2.3E-01 | 26 | 87 |
| Cldn3 | 0.18 | 3.6E-01 | 333 | 100 |
| Cldn12 | -0.10 | 3.9E-01 | 18 | 83 |
| Cldn15 | 0.11 | 4.3E-01 | 109 | 98 |
| Cldn2 | -0.26 | 4.4E-01 | 79 | 98 |
| Tjp1 | -0.11 | 5.0E-01 | 20 | 86 |
| Cldn7 | 0.13 | 5.1E-01 | 309 | 100 |
| Cldn25 | 0.09 | 5.6E-01 | 15 | 79 |
| Ocln | 0.09 | 6.3E-01 | 33 | 90 |
| Myosin |  |  |  |  |
| **Mylk** | **-0.90** | **2.3E-06** | **16** | **80** |
| **Myl9** | **-2.07** | **2.9E-04** | **29** | **89** |
| **Myh11** | **-2.03** | **5.2E-05** | **13** | **76** |
| Myh9 | -0.01 | 9.2E-01 | **32** | **91** |

At the same time, myosin light chain kinase (Mylk) and the myosin II chains Myl9 and Myh11 were downregulated.

Table 3 lists four aquaporin genes with biologically relevant expression. Most notably, the highly abundant water channel aquaporin 8 (Aqp8) is upregulated 375-fold, the second highest absolute log-fold change out of 19,863 genes.

**Table 3:**
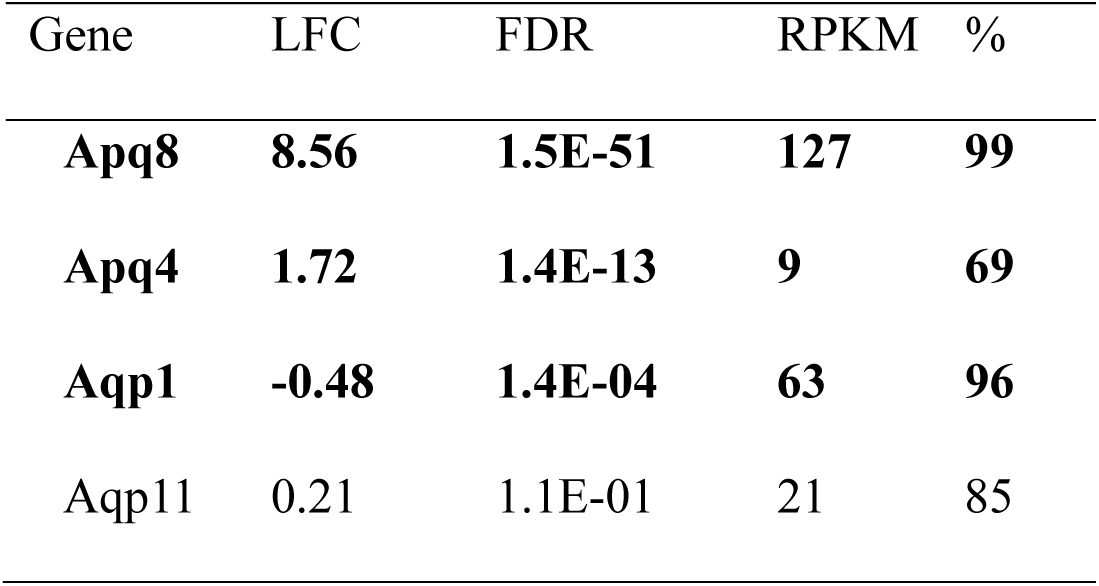
Differential expression and relative abundance of aquaporin water transporter genes. The most abundant aquaporin in humanized gut, Aquaporin 8, was upregulated 375-fold, changing from virtually absent in germ-free colon to highly abundant in humanized microbiome.

### Permeability

Ussing chamber experiments measuring ^14^C-PEG 4000 flux across small intestinal tissue from germ-free and humanized mice were used to assess intestinal permeability. A two-tailed unpaired t-test was used to compare humanized and germ-free samples (Figure 5), showing that permeability was twice as high in germ-free mice as in humanized mice (P = 0.048).

**Figure 5:**
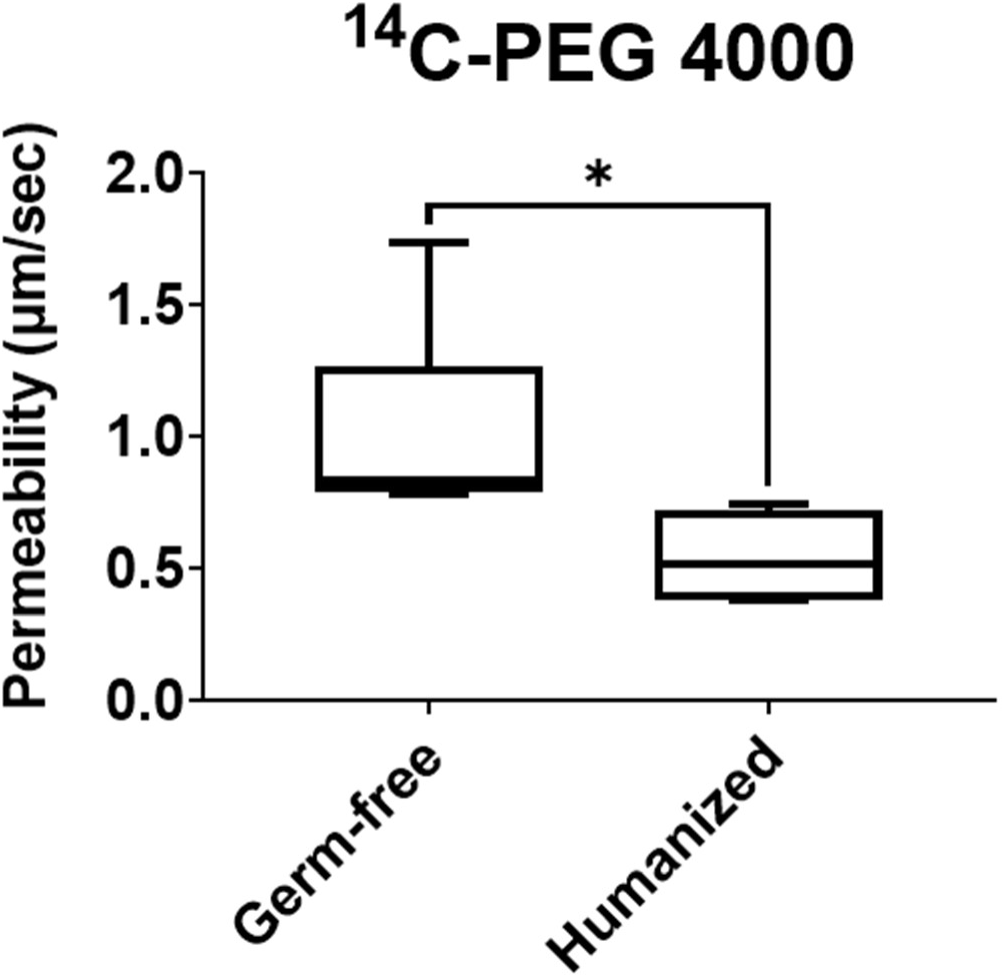
Gut microbiota enhances intestinal barrier integrity: Ussing chamber experiments indicate that small intestinal permeability to ^14^C-PEG 4000 is higher in germ-free mice compared to that of humanized mice (P=0.048).

### Phase 1 drug-metabolizing enzymes

Relative mean expression values for cytochrome P450 genes in germ-free and humanized mouse colons are shown in Figure 6. Out of 20 genes, four are upregulated and three downregulated. None of these, however, correspond to important human drug-metabolizing enzymes. At the same time, cytochrome P450 oxidoreductase (Por), the essential electron donor for all cytochrome P450 activity, was upregulated in humanized colon by 40% (FDR = 9 × 10^-7^).

**Figure 6:**
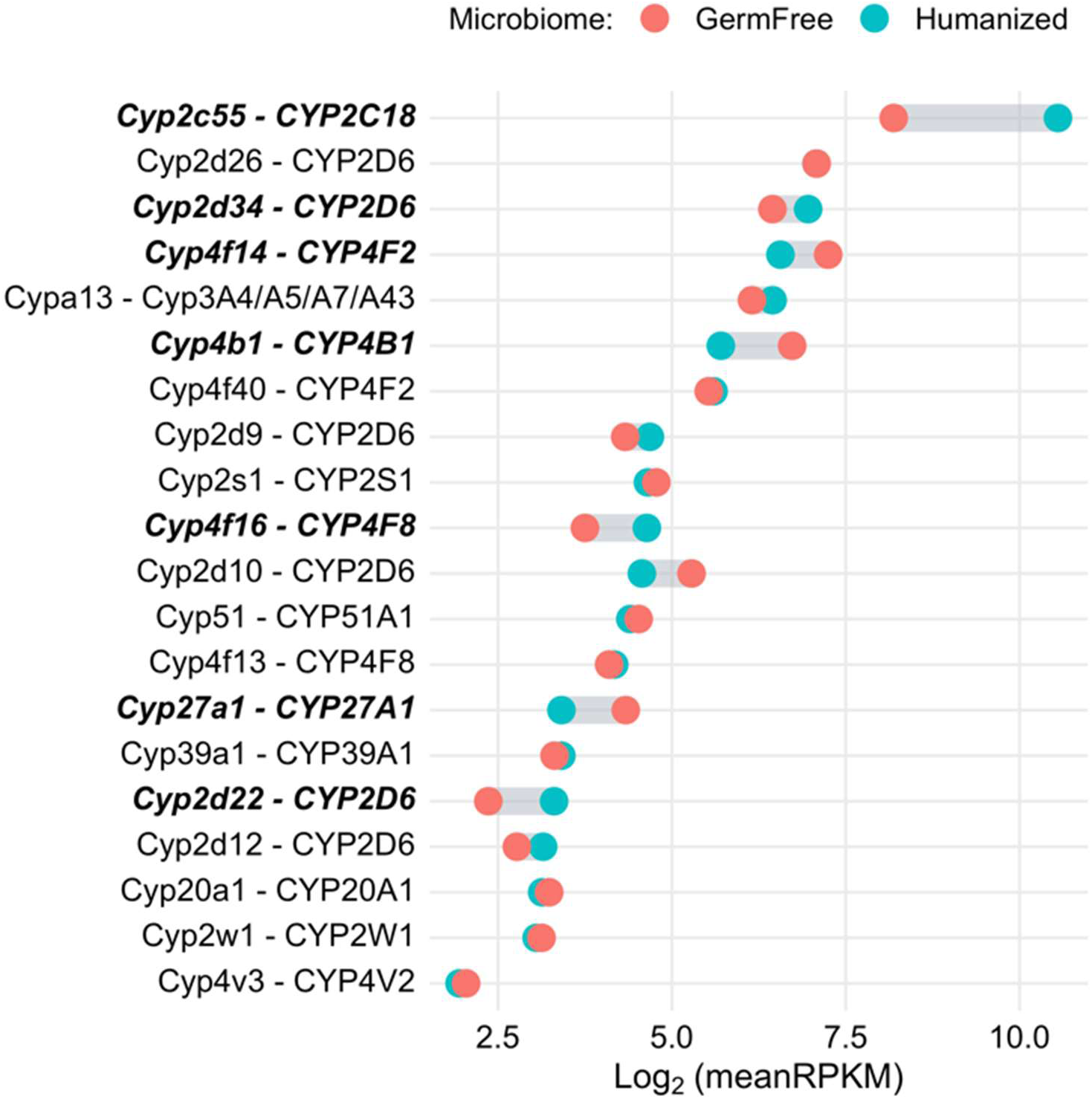
List of cytochrome P450 genes with human homologs and RPKM > 4 (log2(RPKM) > 2). Mus Musculus gene names are given followed by human homologs. Genes in bold are differentially expressed with FDR < 0.01. None of the differentially expressed genes are homologs of human enzymes with known relevance for intestinal drug metabolism.

Differential expression results and relative abundance for a selection of hydrolases and reductases involved in phase 1 metabolism are provided in Table 4. All the mouse homologs of human carboxylesterase 2 (CES2) were upregulated in humanized mice. This includes Ces1a, which increased 24-fold with FDR = 1.3 × 10^-18^. In addition, two isoforms of carbonyl reductase (Cbr1 and Cbr3), the homolog of human aldo-keto reductase AKR1B10, and the quinone reductase Nqo1 were also upregulated.

**Table 4:** Major differentially expressed non-cytochrome-P450 phase 1 enzymes. Included are all carboxylesterse 1, carboxylesterate 2, and carbonyl reductase isoforms with human homologs and RPKM > 4, as well as several other differentially expressed non-cytochrome-P450 reductase enzymes known to have human gut expression. All carboxylesterase 2 homologs are upregulated with very high significance. Carbolylesterase 2 is a major intestinal drug enzyme that hydrolyzes esters, amides, carbamates and thioesters. In humans, only carbolylesterse 2 is expressed in the intestine. Carbonyl reductase reduces carbonyl compounds, including quinones. The aldo/keto reductase AKRIB10 acts on aldehydes and retinoic acid; NQO1 reduces quinones, and cytochrome p450 oxidoreductase (POR) is a critical electron donor for CYP450 activity.

| Gene | Homolog | LFC | FDR | RPKM | % |
| --- | --- | --- | --- | --- | --- |
| Carboxylesterase 2 |  |  |  |  |  |
| **Ces2a** | **CES2** | **4.58** | **1.3E-183** | **137** | **58** |
| **Ces2b** | **CES2** | **2.67** | **1.1E-41** | **26** | **50** |
| **Ces2c** | **CES2** | **1.24** | **1.4E-23** | **248** | **98** |
| **Ces2e** | **CES2** | **1.15** | **1.2E-07** | **189** | **97** |
| Carbonyl reductase |  |  |  |  |  |
| **Cbr3** | **CBR3** | **2.75** | **1.1E-55** | **32** | **54** |
| **Cbr1** | **CBR1** | **1.05** | **3.4E-30** | **169** | **97** |
| Cbr4 | CBR4 | 0.16 | 3.4E-01 | 6 | 56 |
| Carboxylesterase 1 |  |  |  |  |  |
| **Ces1g** | **CES1** | **1.14** | **1.8E-06** | **93** | **93** |
| Ces1d | CES1 | -0.47 | 1.9E-02 | 43 | 95 |
| Other Phase 1 |  |  |  |  |  |
| **Akr1b8** | **AKR1B10/15** | **2.02** | **3.5E-19** | **25** | **60** |
| **Nqo1** | **NQO1** | **0.73** | **1.0E-10** | **39** | **87** |
| **Por** | **POR** | **0.51** | **9.0E-07** | **45** | **90** |

### Phase 2 drug-metabolizing enzymes

Five phase 2 enzyme groups that catalyze acetylation, glucuronidation, glutathione S-conjugation, methylation, and sulfoconjugation were analyzed for over-represented differential expression and the results are presented in Table 5. No phase 2 enzymes were downregulated, while upregulated UDP-glucuronosyltransferases and glutathione S-transferases were significantly over-represented. In addition, all five genes coding enzymes that catalyze the corresponding co-substrates UDP glucuronic acid and glutathione were upregulated, as shown in Table 6.

**Table 5:** Over-representation analysis for major groups of phase II drug metabolizing enzymes. Included are drug metabolizing enzymes with mean RPKM > 4 in either treatment group. Upregulated glutathione S-transferases and UDP–glucuronosyltransferases were over-represented by Fisher’s hypergeometric exact test with p-values of 4.05*10-5 and 2.61*10-4, respectively. UGTs specifically refers to UDP glucuronosyltransferases, which are from the 1A, 2A and 2B families.

| Process | Genes | Total | upreg | Pval(up) | downreg | Pval(dn) |
| --- | --- | --- | --- | --- | --- | --- |
| glucuronidation | UGTs | 12 | 8 | 8.6e-04 | 0 | 1.00 |
| glutathione conj. | GSTs, MGSTs | 18 | 12 | 4.0e-05 | 0 | 1.00 |
| sulfonation | SULTs | 5 | 2 | 0.29 | 0 | 1.00 |
| acetylation | NATs | 2 | 0 | 1.00 | 0 | 1.00 |
| methylation | xxMTs | 3 | 0 | 1.00 | 0 | 1.00 |

**Table 6:** Differential expression results for genes coding enzymes that catalyze co-substrates of UDP–glucuronosyltransferases and glutathione S-transferases: UDP glucuronic acid and glutathione. All genes show statistically significant increases between 30% and 40% (0.40 ≤ LCF ≤ 49).

| Gene | Enzyme | Final co-substate | LFC | FDR |
| --- | --- | --- | --- | --- |
| UGP2 | UDP-glucose pyrophosphorylase | UDP glucuronic acid | 0.43 | 1.0e-04 |
| UGDH | UDP-glucose 6-dehydrogenase | UDP glucuronic acid | 0.40 | 7.7e-03 |
| GCLC | γ-glutamylcysteine synthetase | glutathione | 0.41 | 5.0e-04 |
| GCLM | γ-glutamylcysteine synthetase | glutathione | 0.49 | 1.7e-06 |
| GSS | glutathione synthetase | glutathione | 0.46 | 1.7e-05 |

### Transport: ATP-binding cassettes and solute carriers

The seven most abundant differentially expressed ATP-binding cassette transporters were upregulated in humanized mice, as shown in Figure 7. These include mouse homologs of the human gene ABCB1, which codes for the major drug efflux transporters p-glycoprotein (P-gp): Abcb1a (LFC =0.75, FDR = 8 × 10^-7^) and Abcb1b (LFC = 0.58, FDR = 5 × 10^-3^). Also upregulated were the homologs of multidrug resistance-associated protein 3 (MRP3, encoded by Abcc3) and Abcg2, a homolog of breast cancer resistant protein (BCRP). The mouse gene Abcg3, another BCRP homolog, was expressed at very low levels and not differentially expressed.

**Figure 7:**
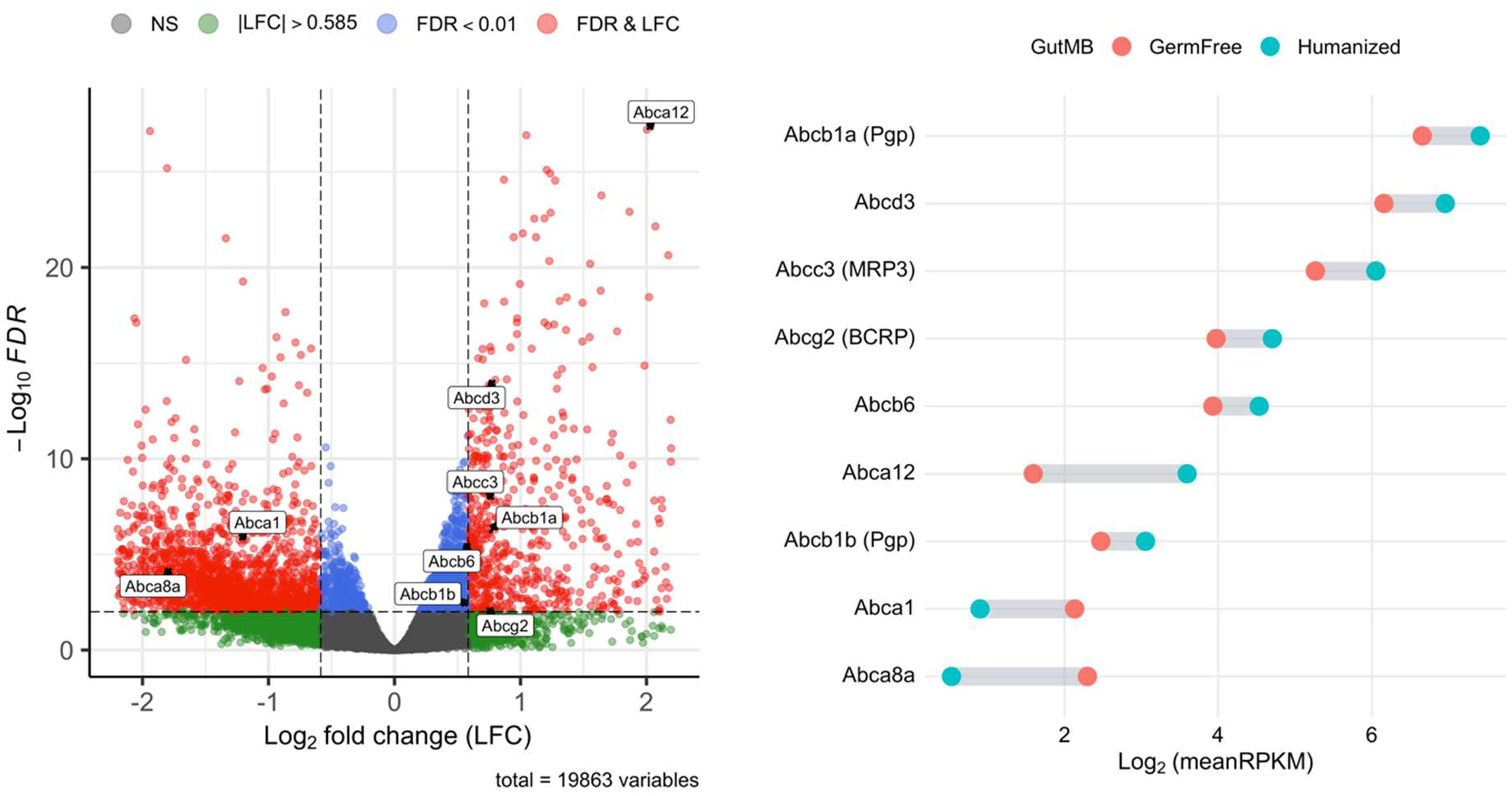
Left, volcano plot of differentially expressed ATP-binding cassette genes with RPKM > 4 (log2(RPKM) > 2). Both mouse homologs of the human gene ABCB1, which codes for the major drug efflux transporters p-glycoprotein (P-gp), are significantly upregulated: Abcb1a (LFC =0.75, FDR = 8*10-7); Abcb1b (LFC = 0.58, FDR = 5*10-3). Also upregulated are the genes Abcc3 (LFC = 0.79, FDR = 1.4*10-8), which codes for multidrug resistance-associated protein 3 (MRP3), and Abcg2 (LCF = 0.73, FDR = 0.006), which codes for breast cancer resistant protein (BCRP). The mouse gene Abcg3, which is also a homolog for ABCG2 (BCRP), was found in very low abundance and not differentially expressed. Right, relative abundance indicated by log2(RPKM). Abcb1a transcripts exhibit very high relative abundance with the greatest abundance of all differentially expressed ATP-binding cassette genes and about 20 times greater than Abcb1b. In humans, ABCA12 is primarily expressed in the skin. Abcb6 and Abcd3 are involved in iron and fatty acid transport, respectively.

An observed increase in P-gp protein expression by immunofluorescence shown in Figure 8 was consistent with transcriptomic results. The mean intensity in humanized mice increased by 77% with a p-value of 0.0012 (C) --roughly equivalent to the 68% upregulation of Abcb1a expression. Additionally, the localization of P-gp on the apical surface was shown to be much higher in humanized mice compared to germ free colon (D).

**Figure 8:**
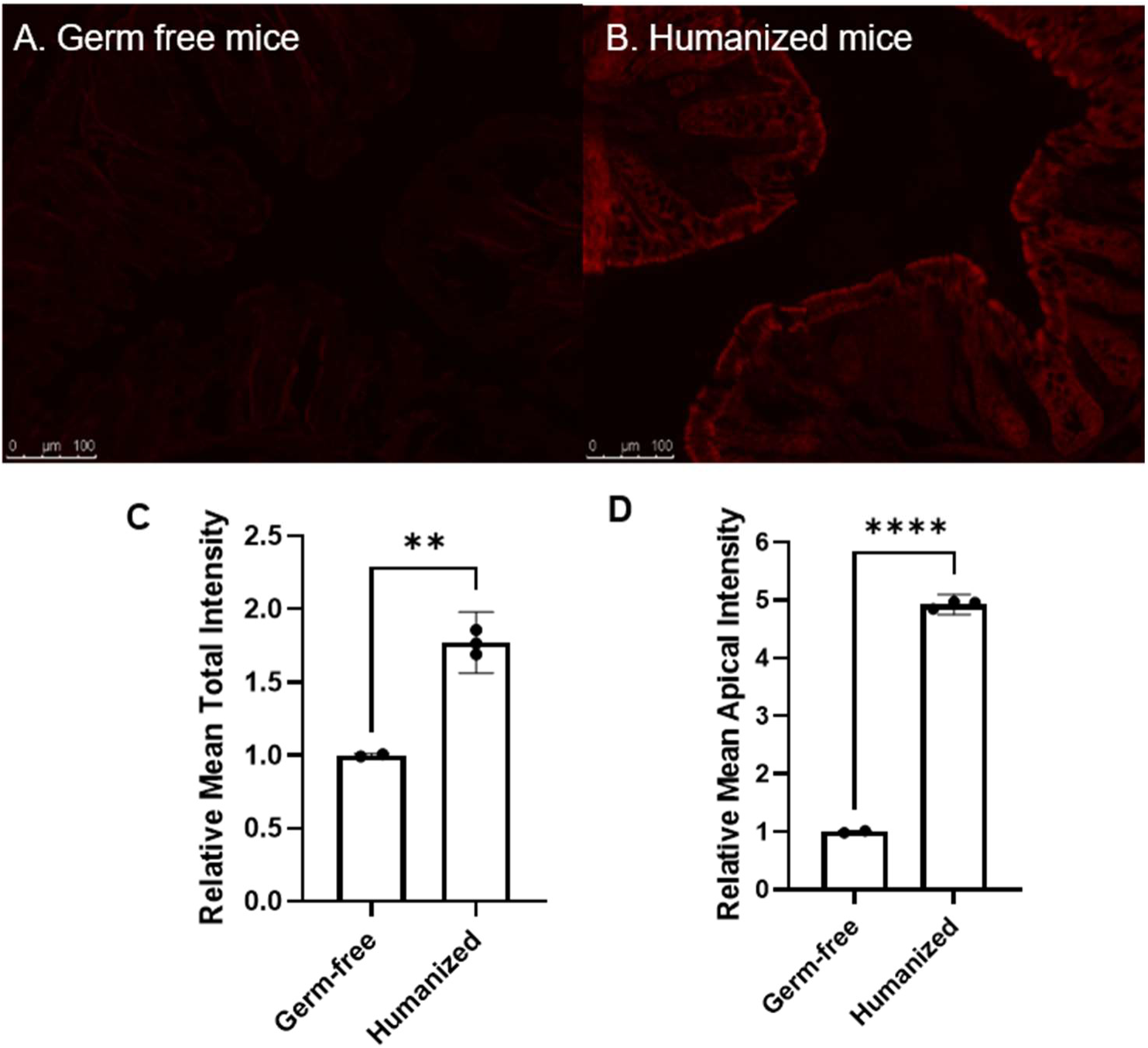
Decreased P-gp expression in the colon of germ-free mice shown by immunohistochemistry. Colon sections from germ-free (A) and humanized (B) mice were stained for P-gp, tagged with Cy3-conjugated streptavidin dye. (C) Relative mean intensity of two germ-free and three humanized mice with 95% confidence intervals, along with mean intensity along apical surface shown in (D). Dots represent Scale bar: 100 μm. Fold-changes were 1.77 and 4.95, respectively. In unpaired t-test, p-values were, 0.0012 and < 0.0001 (**p < 0.01, ****p < 0.0001).

Table 7 provides a full list of differentially expressed solute carriers with known drug substrates. Notably, the major apical influx intestinal drug transporters--monocarboxylate transporter 1 (MCT1), encoded by Slc16A1, and organic cation transporter novel family member 2 (OCTN2), encoded by Slc22A5--were upregulated.

**Table 7:** Differentially expressed solute carriers with known human homologs and drug substrates filtered by FDR < 0.01 and RPKM > 4. Shown in bold are the major apical influx intestinal drug transporters Slc16a1, which codes for the monocarboxylate transporter 1 (MCT1), and SLC22a5, which codes the organic cation transporter novel family member 2 (OCTN2), are both upregulated in humanized colon.

| Gene | LFC | FDR | RPKM | Endogenous | Drug Substrates |
| --- | --- | --- | --- | --- | --- |
| Slc25a37 | -0.74 | 3.7E-16 | 21 | Metals | Mercurials, Tannic Acid |
| **Slc16a1** | **1.00** | **1.5E-10** | **28** | **Carboxylates** | **Anticancer prodrugs, NSAIDs** |
| Slc39a14 | -0.51 | 2.4E-10 | 7 | Metals | Antidepressants |
| Slc23a2 | 0.57 | 1.2E-07 | 10 | Carboxylates | Vitamin C |
| Slc40a1 | 1.09 | 4.3E-07 | 23 | Metals | CNS Stimulants, Hormones |
| Slc5a8 | 0.82 | 3.0E-06 | 36 | Fatty Acids | NSAIDs |
| Slc25a10 | 0.59 | 3.7E-06 | 55 | Carboxylates | Succinic Acid |
| Slc13a1 | -0.72 | 1.7E-05 | 21 | Inorganic Ions | Thiosulfate, Selenate, Molybdate, Citrate, Succinic Acid |
| Slc9a3 | 0.72 | 1.5E-04 | 31 | Metals | Cariporide |
| Slc25a11 | 0.28 | 2.0E-04 | 35 | Carboxylates | Oxogluric Acid |
| Slc25a5 | 0.41 | 4.4E-04 | 423 | Glycosylamines | Clodronate |
| **Slc22a5** | **0.39** | **4.7E-04** | **21** | **Organic Ions** | **Valproic Acid, Verapamil, Imatinib** |
| Slc25a22 | 0.32 | 5.3E-04 | 8 | Amino Acids | Glutamic Acid |
| Slc25a3 | 0.34 | 6.6E-04 | 98 | Inorganic Ions | Phosphate |
| Slc10a2 | -0.67 | 7.1E-04 | 207 | Bile Acids | Anticancer Agents, Antiviral Agents |
| Slc31a1 | 0.43 | 9.6E-04 | 23 | Metals | Anticancer Agents |
| Slc38a1 | 0.39 | 1.3E-03 | 6 | Amino Acids | Lithium |
| Slc25a34 | 0.45 | 1.6E-03 | 15 | Glycosylamines | Mercurials, Tannic Acid |
| Slc25a13 | 0.30 | 2.9E-03 | 14 | Amino Acids | Calcium |
| Slc25a24 | 0.38 | 6.8E-03 | 97 | Cofactors | Calcium |

### Efflux and cell viability studies (organoids)

Live imaging of colonic organoids incubated with calcein-AM revealed differences in fluorescence intensity between experimental groups. In organoids from both germ-free and humanized colon, cyclosporin A pretreatment increased intracellular fluorescence along the apical surface, compared to untreated controls, suggesting reduced calcein-AM efflux by P-gP. In addition to apical fluorescence differences with and without inhibitor treatment, organoids from humanized mice colon exhibited higher fluorescence intensity than those from germ-free mice.

This may reflect increased hydrolysis of calcein-AM by esterases, particularly carboxylesterase 2 (CES2), which was significantly upregulated in humanized mice (Table 4). Representative images for each condition are shown in Figure 9.

**Figure 9:**
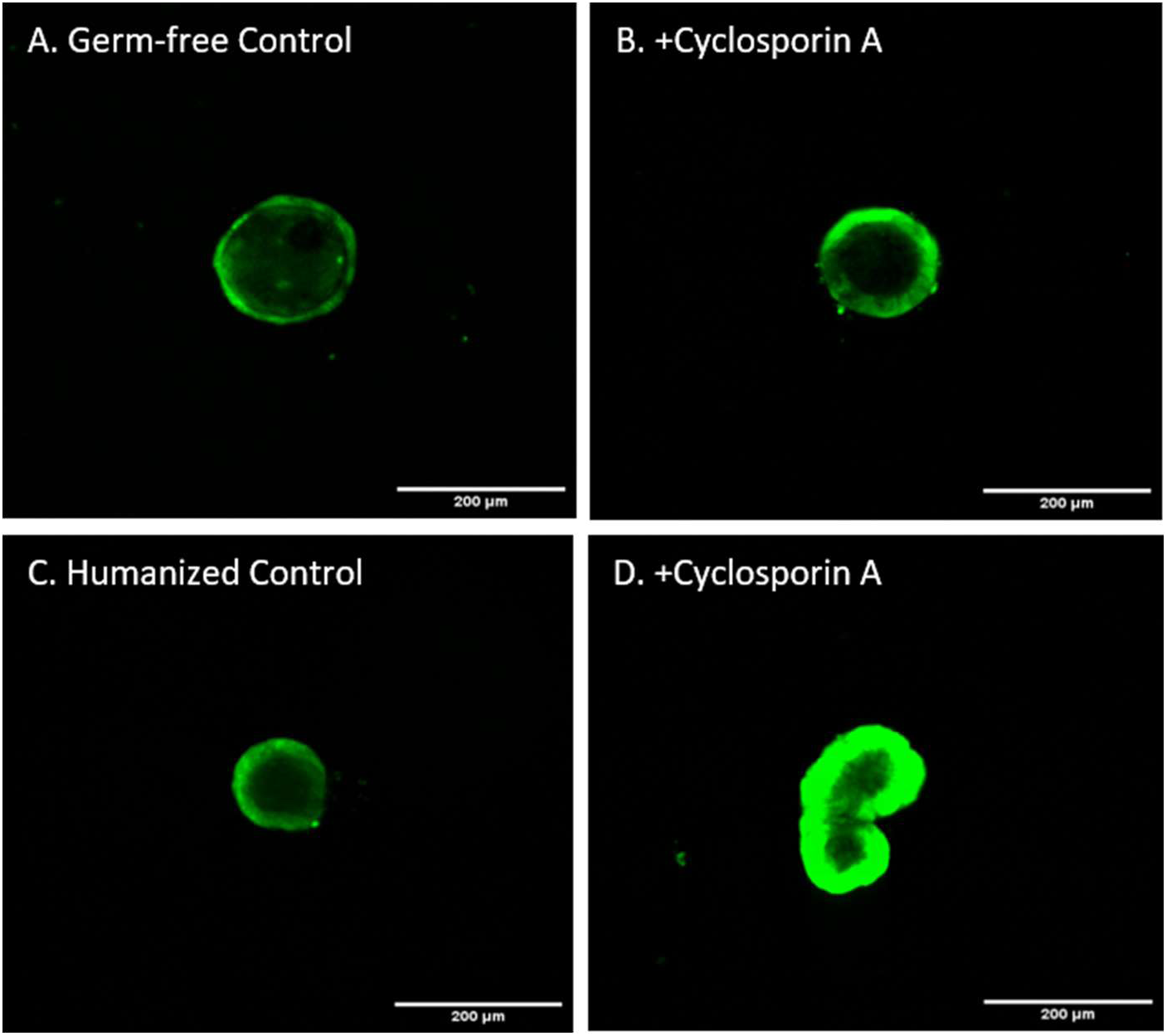
Efflux transporter, P-gp functional study: Colonoids from germ-free and humanized mice were pre-treated with P-gp inhibitor (Cyclosporin A) for 30 minutes followed by treatment with Calcein-AM for 60 min at room temperature. Confocal images show Calcein-AM accumulation in germ-free colonid control (A), and cyclosporine A pre-treated colonoid (B) along with control from humanized mice (C) and cyclosporine A pre-treated colonoid (D). Scale bar, 200 μm.

## DISCUSSION

Gut bacteria exert a profound influence on the expression of genes that affect oral drug bioavailability, as demonstrated by the significant differences observed in mouse gut maintained under germ-free conditions compared to that inoculated with humanized microbiota.

Understanding how gut bacteria generally shape intestinal phenotype provides a foundation for characterizing how microbiota variability, particularly gut dysbiosis, may impact oral drug bioavailability.

In this study, the humanized mouse cohort was colonized with the same human fecal sample to ensure consistent microbiota composition across all humanized mice. While this intra-group uniformity does not reflect the natural diversity of human microbiota, it enabled confident and comprehensive identification of bacteria-sensitive genes using a limited sample size. The effectiveness of this approach is illustrated by well-separated clusters obtained from PCA. After employing a strict 1% false discovery rate threshold, we identified roughly 20% of 20,000 microbiota-sensitive genes. Among these are many genes with diverse roles in drug disposition: active transporters, drug-metabolizing enzymes, and tight junctions regulating paracellular passive diffusion. Also included are other factors with less direct influence related to immune responses, extracellular matrix turnover, and water transport. The PCA plot also revealed greater variability in gene expression among humanized mice compared to germ-free mice. This highlights microbiota as a source of phenotypic variability, which likely involves some degree of microbiota heterogeneity developed eight weeks following the introduction of a homogeneous fecal sample and variability in host response. While not included in this study, a metagenomic analysis could clarify the extent of this compositional drift, its difference from conventional mouse microbiota, and its contribution to phenotypic variability.

As expected, genes related to both innate and adaptive immune responses were upregulated in humanized mice, but two distinct patterns emerged. Genes involved in innate immune responses and nonspecific processes, to such as NOS2 (inducible nitric oxide synthase), components of the dual oxidase system (DUOX1, DUOX2, DUOXA1, DUOXA2), and PIGR (polymeric immunoglobulin receptor), which facilitates IgA transport to the mucosa, exhibited uniformly low dispersion across samples. This consistency resulted in low p-values relative to all other genes, despite moderate fold-changes. Expression of NOS2 and DUOX2 has been shown to be inducible by lipopolysaccharides, which are components of gram-negative bacterial membranes.^17^ In contrast, genes involved in adaptive immune responses, including various constant and variable immunoglobulin chains that form IgA and IgG antibodies, were among the most upregulated, but also exhibited very high variability, reflected in their relatively high p-values. While antibodies alter mucus production and can bind biologics, the direct effects of immune response on oral drug disposition are limited. The most relevant consideration in the context of this study is the bidirectional influence of immune response and microbiota composition^18,19^ as a potential source of individual phenotypic variability.

In this study, genes encoding constituents and regulators of the ECM were found to be generally attenuated in the presence of gut bacteria, including all the major collagen and laminin isoforms that form the gut epithelial basement membrane. The ECM regulates the integrity of the gut epithelium and is itself a regulator of gene expression via both ligand-receptor signaling and mechanotransduction.^20,21^ While the ECM itself does not present a barrier to drug transport, pathological increases in MMP protease activity--particularly MMP2 and MMP9, which were upregulated along with MMP7 in germ-free mice--are linked to dysregulated ECM turnover and loss of tight junction integrity.^21^

Ussing chamber permeability of PEG 4000 through the small intestinal mucosal explant was higher in germ-free mice compared to mice with human microbiota. To investigate possible reasons for this difference, we analyzed genes related to tight junction structure (claudins, occludin, and zonula occludens proteins) and actomyosin contractility (myosin II and myosin light chain kinase, regulating myosin phosphorylation). Although transcriptomic analysis was limited to the colon where the presence of bacterial populations are several magnitudes higher compared to that in the small intestine, the strong and compelling results obtained in the colon would allow us to speculate that similar changes are possible in the small intestine as well.

Two distinct pathways mediate tight junction permeability: the pore pathway, which facilitates high-throughput transport with specific size and charge selectivity, and the leak pathway, a more general route for larger solutes such as PEG 4000. The leak pathway is primarily regulated by proteins such as zonula occludens and occludin, while selective pores are formed by certain claudin proteins.^22^ Claudin-4 has been shown to both form pores and decrease leak-pathway transport, possibly through indirect mechanisms.^22^

In humanized mice colon, Cldn4 and Tjp3 (encoding ZO-3) were upregulated, suggesting greater tight junction integrity.^23,24^ Additionally, reduced expression of Mlck (myosin light chain kinase) and genes coding myosin II heavy and light chains suggest a less dynamic, more stable actomyosin cytoskeleton.^25^ Various studies have demonstrated that pathological phosphorylation of myosin light chain by MLCK leads to greater permeability.^26,25^

The most significant differences were the higher humanized expression of Cldn4 and Cldn23. Interestingly, recent studies show that CLDN4 and CLDN23 form paracellular channels together with uniquely high size and charge selectivity, and in some orientation may form poreless junctions.^27^ This suggests a mechanism by which microbiota may also restrict paracellular movement through the pore pathway.

Among the most dramatic changes induced by microbiota is the 375-fold upregulation of Aqp8, a homolog of the major colonic water channel Aquaporin 8.^28^ Based on conditions in other tissues that stimulate Aquaporin expression, two factors likely drive the increased expression of this transporter: an osmotic gradient^29,30^ from epithelial absorption of bacterial fermentation metabolites and H2O2 production^31^ by dual oxidase enzymes. The expected result is decrease in luminal water content^28^; alleviating diarrhea found in germ-free mice. This could impact bioavailability of low-permeable drugs by affecting water content required for drug dissolution and transit time.

Our findings did not support any relevant microbiota influence on cytochrome P450 gene expression. As shown in Figure 6, there were roughly equal numbers of upregulated and downregulated Cyp genes, none of which are associated with human drug metabolism. The main human phase I drug-metabolizing enzymes are members of the CYP1, CYP2, or CYP3 cytochrome P450 family.^32^ In both the liver and the intestine, CYP3A is the predominant subfamily in activity and content, particularly CYP3A4.^32^ The CYP3A family accounts for roughly one-third of cytochrome P450 enzyme abundance in the liver, while it comprises more than two-thirds in the intestine.^32^ The intestinal drug-metabolizing CYP3A enzymes--CYP3A4, CYP3A5, CYP3A7, and CYP3A43,--are all homologs of the mouse cytochrome Cyp3a13. As expected, Cyp3a13 was found in significant abundance, although it was not differentially expressed. Of secondary importance in the human intestine is the CYP2C family^32^, of which only the homolog of CYP2C18 (Cyp2c55) was found. Of the known intestinal drug-metabolizing CYP2C enzymes, CYP2C18 appears to have the lowest protein abundance or clinical importance, although mRNA content has been found to be significant. Among other homologs of human intestinal drug metabolizing enzymes with possible minor significance^33^ – including CYP2J2, CYP2D6, CYP1A1, CYP2E1, and CYP2S1 – only CYP2S1 was detected, and it was also not differentially expressed.

The lack of bacterial influence on cytochrome P450 expression may still be considered inconclusive in humans, as cytochrome metabolism of drugs in mice is divergent and generally more efficient, employing a distinct and much larger set of isoforms.^34^ This point is illustrated by the fact that much more accurate drug pharmacokinetic mouse models have recently been developed by replacing 33 genes from drug-metabolizing Cyp1a, 2c, 2d, and 3a families with six key human enzymes: CYP1A1, CYP1A2, CYP2C9, CYP2D6, CYP3A4, and CYP3A7.^35,36^

In contrast to the lack of microbiota-induced expression of cytochrome P450 oxidizers, expression of several non-cytochrome phase I hydrolase and reductase genes exhibited significant, in some cases profound, stimulation in humanized mice. After cytochrome oxidation, hydrolysis is the second most common phase I drug-metabolizing process and has been the most common enzymatic reaction for the activation of prodrugs.^37^ The most important human drug-metabolizing intestinal hydrolase is Carboxylesterase 2, which is involved in hydrolyzing esters to carboxylic acids in compounds such as Clopidogrel and the prodrug Oseltamivir.^38,39^ Four Carboxylesterase 2 homologs (Ces2a, Ces2b, Ces2c and Ces2e) were expressed, all of which were substantially upregulated. Ces2a was increased 24-fold in humanized mice with an FDR on the order to 10^-183^. Carboxylesterase 1 was also upregulated. In humans, however, this is the predominant liver isoform without significant expression in the gut.^38^ Upregulated reductases with relevant human intestinal activity include homologs of Carbonyl reductase 1 (Cbr1), Aldo-Keto Reductase Family 1 Member B10 (Akr1b8), NAD(P)H Quinone Dehydrogenase 1 (Nqo1), and NADPH-Cytochrome P450 Reductase (Por). Human AKR1B10 and NQO1 are important detoxifying enzymes, reducing aldehydes, ketones, and quinones.^40–42^ POR, on the other hand, is a ubiquitous enzyme and essential electron doner for cytochrome P450 function.^43^ While the results do not support gut bacteria influence on cytochrome expression, the upregulation of Por suggests an increased potential for cytochrome activity.

Table 5 demonstrated that upregulated glutathione S-transferases and UDP-glucuronosyltransferases were statistically over-represented. The latter represents the weightiest of phase II drug-metabolizing enzymes with a broad substrate panel that includes non-steroidal anti-inflammatory agents.^37,44^ Increased activity of both these phase II processes is also supported by the upregulation of the enzymes that synthesize the corresponding co-substrates: UDP-glucuronic acid and glutathione, as shown in Table 6.

Eight of the 12 UDP-glucuronosyltransferase genes were found to be upregulated with FDR less than 0.01. This includes all four expressed mouse homologs (Ugt1a7c, Ugt1a8, Ugt1a9, and Ugt1a10) that map to UGT1A10, the most abundant A1-UDP glucuronosyltransferase in both colon and intestine.^44^ On the other hand, 2B enzymes were not differentially expressed. Despite the clear upregulation of 1A-UDP glucuronosyltransferases along with co-substrate-producing enzymes, the difference in potential glucuronidation of drugs is confounded by the large and active concentration of microbial b-glucuronidase enzymes, which reverse glucuronidation and supply glucuronic acid as a bacterial energy substrate.^45^ While β-glucuronidase activity likely plays a major role in the observed expression changes, the degree to which the microbiota-mediated increase in UDP glucuronosyltransferase expression and production of UDP glucuronic acid compensates for the activity of b-glucuronidases is unknown.

Microbiota metabolism is likely also a stimulator of glutathione S-transferase expression, as these enzymes have been shown to be induced by products of gut fermentation, particularly butyrate.^46^ In addition to drug detoxification, glutathione S-transferases are particularly important for the detoxification of reactive microbial metabolites.^47^

Closely linked with phase II conjugation is active transport by ATP-binding cassettes drug transporters, as soluble glucuronides and glutathione conjugates are substrates of these transporters.^48^ Arguably, the most clinically relevant finding is the increased expression of P-gp. This predominant drug efflux transporter reduces bioavailability for a wide range of therapeutics, particularly moderately lipophilic, neutral, or weakly basic drugs, with limited hydrogen bonding, allowing lipid bilayer entry before efflux.^1,49^ Mean immunofluorescence measurements suggested a ∼2-fold increase in P-gp expression, consistent with the magnitude of gene upregulation. Homologs of efflux transporters, BCRP and multidrug-resistance associated protein 3 (MRP3), also showed gene upregulation in humanized colon. Both P-pg and BCRP activity in the enterocytes decrease bioavailability, while MRP3 is localized at the basolateral surface and may actually increase bioavailability of drug substrates or enzyme-activated prodrugs.^50^ Linked to phase II metabolism, MRP3 transports glutathione, sulfate or glucuronide conjugates of substrates such as etoposide and methotrexate.^51^ Thus, it’s upregulation may be linked to our findings on microbiota-mediated changes to phase II enzyme expression.

Among the upregulated solute carriers, SLC22A5 (OCTN2) is considered a significant drug transporter in the human small intestine. It is an apically polarized influx transporter specialized in the absorption of endogenous zwitterion carnitine, a fatty acid shuttle and regulator of acyl-CoA balance.^48^ OCTN2 also functions as an important transporter for many other ionic and neutral compounds, including imatinib (anti-cancer agent), ipratropium (bronchodilator), verapamil (calcium channel blocker), valproic Acid (anticonvulsant/mood stabilizer), and quinidine (antiarrhythmic agent).^9,48^ The homolog human monocarboxylate transporter 1 (MCT1), Slc16a1, demonstrated a two-fold upregulation. Intestinal MCT1 is mainly known for transporting metabolic monocarboxylates such as lactate, pyruvate, and ketone bodies^52^, but is also known as a transporter of weakly acidic drugs with a carboxylic acid moiety, such as Fluorouracil and Gemcitabine prodrugs^53,54^ (anticancer agents), salicylic acid^55^ (NSAID), and γ- Hydroxybutyrate^56^ (narcolepsy treatment).

This study demonstrated microbiota influence on intestinal phenotype in ways that affect oral drug bioavailability and provides a methodological foundation to conduct targeted studies to evaluate the impact of gut microbiome in health and disease on drug disposition of orally administered drugs. In the current study, phenotypic changes in the colon have been characterized. However, it is essential to extend approach to significant sites of drug absorption in the small intestine. Additionally, characterization of the expression of other enzymes and transporters in the small intestine will enable us to link drug disposition with molecular changes triggered by gut microbiota.

## Supporting information

Supplementary file 1: Raw counts

Supplementary file 3: Relative abundance

Supplementary file 2: Differential expression

## REFERENCES

1. Andersen, V. C. & Sonne, J. [Drug metabolism in the small intestine--the significance for biological availability]. Ugeskr Laeger 162, 3215–3219 (2000).

2. Raffa, R. B., Pergolizzi, J. V., Taylor, R., Decker, J. F. & Patrick, J. T. Acetaminophen (Paracetamol) Oral Absorption and Clinical Influences. Pain Practice 14, 668–677 (2014).

3. Gong, L., Goswami, S., Giacomini, K. M., Altman, R. B. & Klein, T. E. Metformin pathways: pharmacokinetics and pharmacodynamics. Pharmacogenet Genomics 22, 820–827 (2012).

4. Marok, F. Z. et al. A Physiologically Based Pharmacokinetic Model of Ketoconazole and Its Metabolites as Drug–Drug Interaction Perpetrators. Pharmaceutics 15, 679 (2023).

5. Pappenheimer, J. R. Paracellular intestinal absorption of glucose, creatinine, and mannitol in normal animals: relation to body size. American Journal of Physiology-Gastrointestinal and Liver Physiology 259, G290–G299 (1990).

6. McDonnell, A. M. & Dang, C. H. Basic review of the cytochrome p450 system. J Adv Pract Oncol 4, 263–268 (2013).

7. Sekirov, I., Russell, S. L., Antunes, L. C. M. & Finlay, B. B. Gut Microbiota in Health and Disease. Physiological Reviews 90, 859–904 (2010).

8. Zhang, X., Han, Y., Huang, W., Jin, M. & Gao, Z. The influence of the gut microbiota on the bioavailability of oral drugs. Acta Pharmaceutica Sinica B 11, 1789–1812 (2021).

9. Estudante, M., Morais, J. G., Soveral, G. & Benet, L. Z. Intestinal drug transporters: An overview. Advanced Drug Delivery Reviews 65, 1340–1356 (2013).

10. Azman, M., Sabri, A. H., Anjani, Q. K., Mustaffa, M. F. & Hamid, K. A. Intestinal Absorption Study: Challenges and Absorption Enhancement Strategies in Improving Oral Drug Delivery. Pharmaceuticals (Basel) 15, 975 (2022).

11. Lemmens, G., Van Camp, A., Kourula, S., Vanuytsel, T. & Augustijns, P. Drug Disposition in the Lower Gastrointestinal Tract: Targeting and Monitoring. Pharmaceutics 13, 161 (2021).

12. Kalari, K. R. et al. MAP-RSeq: Mayo Analysis Pipeline for RNA sequencing. BMC Bioinformatics 15, 224 (2014).

13. Liao, Y., Smyth, G. K. & Shi, W. featureCounts: an efficient general purpose program for assigning sequence reads to genomic features. Bioinformatics 30, 923–930 (2014).

14. Bhattarai, Y. et al. Role of gut microbiota in regulating gastrointestinal dysfunction and motor symptoms in a mouse model of Parkinson’s disease. Gut Microbes 13, 1866974 (2021).

15. Bhattarai, Y. et al. Gut Microbiota-Produced Tryptamine Activates an Epithelial G-Protein-Coupled Receptor to Increase Colonic Secretion. Cell Host & Microbe 23, 775–785.e5 (2018).

16. Pompili, S., Latella, G., Gaudio, E., Sferra, R. & Vetuschi, A. The Charming World of the Extracellular Matrix: A Dynamic and Protective Network of the Intestinal Wall. Front. Med. 8, 610189 (2021).

17. Sommer, F. & Bäckhed, F. The gut microbiota engages different signaling pathways to induce Duox2 expression in the ileum and colon epithelium. Mucosal Immunology 8, 372–379 (2015).

18. Gu, M. et al. Host innate and adaptive immunity shapes the gut microbiota biogeography. Microbiology and Immunology 66, 330–341 (2022).

19. Wu, H.-J. & Wu, E. The role of gut microbiota in immune homeostasis and autoimmunity. Gut Microbes 3, 4–14 (2012).

20. Kim, S.-H., Turnbull, J. & Guimond, S. Extracellular matrix and cell signalling: the dynamic cooperation of integrin, proteoglycan and growth factor receptor. Journal of Endocrinology 209, 139–151 (2011).

21. Vilardi, A., Przyborski, S., Mobbs, C., Rufini, A. & Tufarelli, C. Current understanding of the interplay between extracellular matrix remodelling and gut permeability in health and disease. Cell Death Discov. 10, 258 (2024).

22. Shen, L., Weber, C. R., Raleigh, D. R., Yu, D. & Turner, J. R. Tight Junction Pore and Leak Pathways: A Dynamic Duo. Annu. Rev. Physiol. 73, 283–309 (2011).

23. Lu, Z., Ding, L., Lu, Q. & Chen, Y.-H. Claudins in intestines: Distribution and functional significance in health and diseases. Tissue Barriers 1, e24978 (2013).

24. Ma, T. Y., Nighot, P. & Al-Sadi, R. Tight Junctions and the Intestinal Barrier. in Physiology of the Gastrointestinal Tract 587–639 (Elsevier, 2018). doi:10.1016/B978-0-12-809954-4.00025-6.

25. He, W.-Q. et al. Contributions of Myosin Light Chain Kinase to Regulation of Epithelial Paracellular Permeability and Mucosal Homeostasis. IJMS 21, 993 (2020).

26. Cunningham, K. E. & Turner, J. R. Myosin light chain kinase: pulling the strings of epithelial tight junction function. Annals of the New York Academy of Sciences 1258, 34–42 (2012).

27. Raya-Sandino, A. et al. Claudin-23 reshapes epithelial tight junction architecture to regulate barrier function. Nat Commun 14, 6214 (2023).

28. Liao, S., Gan, L., Lv, L. & Mei, Z. The regulatory roles of aquaporins in the digestive system. Genes & Diseases 8, 250–258 (2021).

29. Schnabel, B., Kuhrt, H., Wiedemann, P., Bringmann, A. & Hollborn, M. Osmotic regulation of aquaporin-8 expression in retinal pigment epithelial cells in vitro: Dependence on KATP channel activation. Mol Vis 26, 797–817 (2020).

30. Lin, C. et al. Osmotic pressure induces translocation of aquaporin-8 by P38 and JNK MAPK signaling pathways in patients with functional constipation. Digestive and Liver Disease 55, 1049–1059 (2023).

31. Liu, S.-H. et al. Aquaporin-8 promotes human dermal fibroblasts to counteract hydrogen peroxide-induced oxidative damage: A novel target for management of skin aging. Open Life Sciences 19, 20220828 (2024).

32. Zhao, M. et al. Cytochrome P450 Enzymes and Drug Metabolism in Humans. IJMS 22, 12808 (2021).

33. Dressman, J. B. & Thelen, K. Cytochrome P450-mediated metabolism in the human gut wall. j pharm pharmacol 61, 541–558 (2009).

34. Uehara, S. et al. Comparison of mouse and human cytochrome P450 mediated-drug metabolising activities in hepatic and extrahepatic microsomes. Xenobiotica 52, 229–239 (2022).

35. MacLeod, A. K. et al. Acceleration of infectious disease drug discovery and development using a humanized model of drug metabolism. Proc. Natl. Acad. Sci. U.S.A. 121, e2315069121 (2024).

36. Le Bras, A. Humanized mouse models of drug metabolism. Lab Anim 53, 87–87 (2024).

37. Cerny, M. A. Prevalence of Non–Cytochrome P450–Mediated Metabolism in Food and Drug Administration–Approved Oral and Intravenous Drugs: 2006–2015. Drug Metabolism and Disposition 44, 1246–1252 (2016).

38. Imai, T. & Ohura, K. The Role of Intestinal Carboxylesterase in the Oral Absorption of Prodrugs. CDM 11, 793–805 (2010).

39. Wang, D. et al. Human carboxylesterases: a comprehensive review. Acta Pharmaceutica Sinica B 8, 699–712 (2018).

40. Fukami, T., Yokoi, T. & Nakajima, M. Non-P450 Drug-Metabolizing Enzymes: Contribution to Drug Disposition, Toxicity, and Development. Annu. Rev. Pharmacol. Toxicol. 62, 405–425 (2022).

41. Hao, H. et al. Identification of a Novel Intestinal First Pass Metabolic Pathway: NQO1 Mediated Quinone Reduction and Subsequent Glucuronidation. CDM 8, 137–149 (2007).

42. Endo, S., Matsunaga, T. & Nishinaka, T. The Role of AKR1B10 in Physiology and Pathophysiology. Metabolites 11, 332 (2021).

43. Hamdane, D. et al. Structure and Function of an NADPH-Cytochrome P450 Oxidoreductase in an Open Conformation Capable of Reducing Cytochrome P450. Journal of Biological Chemistry 284, 11374– 11384 (2009).

44. Rowland, A., Miners, J. O. & Mackenzie, P. I. The UDP-glucuronosyltransferases: Their role in drug metabolism and detoxification. The International Journal of Biochemistry & Cell Biology 45, 1121–1132 (2013).

45. Pellock, S. J. & Redinbo, M. R. Glucuronides in the gut: Sugar-driven symbioses between microbe and host. Journal of Biological Chemistry 292, 8569–8576 (2017).

46. Ebert, M. N. Expression of glutathione S-transferases (GSTs) in human colon cells and inducibility of GSTM2 by butyrate. Carcinogenesis 24, 1637–1644 (2003).

47. Peters, W. H. M., Roelofs, H. M. J., Nagengast, F. M. & Van Tongeren, J. H. M. Human intestinal glutathione *S*-transferases. Biochemical Journal 257, 471–476 (1989).

48. El-Kattan, A. & Varm, M. Oral Absorption, Intestinal Metabolism and Human Oral Bioavailability. in Topics on Drug Metabolism (ed. Paxton, J.) (InTech, 2012). doi:10.5772/31087.

49. Li, D. et al. ADMET Evaluation in Drug Discovery. 13. Development of *in Silico* Prediction Models for P-Glycoprotein Substrates. Mol. Pharmaceutics 11, 716–726 (2014).

50. Murakami and Takano - 2008 - Intestinal efflux transporters and drug absorption.pdf.

51. Borst, P., De Wolf, C. & Van De Wetering, K. Multidrug resistance-associated proteins 3, 4, and 5. Pflugers Arch - Eur J Physiol 453, 661–673 (2007).

52. Halestrap, A. P. Monocarboxylic Acid Transport. in *Comprehensive Physiology* (ed. Terjung, R.) 1611–1643 (Wiley, 2013). doi:10.1002/cphy.c130008.

53. Sun, Y. et al. A novel oral prodrug-targeting transporter MCT 1: 5-fluorouracil-dicarboxylate monoester conjugates. Asian Journal of Pharmaceutical Sciences 14, 631–639 (2019).

54. Wang, Y. et al. A facile di-acid mono-amidation strategy to prepare cyclization-activating mono-carboxylate transporter 1-targeting gemcitabine prodrugs for enhanced oral delivery. International Journal of Pharmaceutics 573, 118718 (2020).

55. Watanabe, H., Yashiro, T., Tohjo, Y. & Konishi, Y. Non-Involvement of the Human Monocarboxylic Acid Transporter 1 (MCT1) in the Transport of Phenolic Acid. Bioscience, Biotechnology, and Biochemistry 70, 1928–1933 (2006).

56. Lam, W. K., Felmlee, M. A. & Morris, M. E. Monocarboxylate Transporter-Mediated Transport of γ- Hydroxybutyric Acid in Human Intestinal Caco-2 Cells. Drug Metabolism and Disposition 38, 441–447 (2010).

